# RobNorm: Model-Based Robust Normalization Method for Labeled Quantitative Mass Spectrometry Proteomics Data

**DOI:** 10.1101/770115

**Authors:** Meng Wang, Lihua Jiang, Ruiqi Jian, Joanne Y. Chan, Qing Liu, Michael P. Snyder, Hua Tang

## Abstract

**Motivation:** Data normalization is an important step in processing proteomics data generated in mass spectrometry (MS) experiments, which aims to reduce sample-level variation and facilitate comparisons of samples. Previously published methods for normalization primarily depend on the assumption that the distribution of protein expression is similar across all samples. However, this assumption fails when the protein expression data is generated from heterogenous samples, such as from various tissue types. This led us to develop a novel data-driven method for improved normalization to correct the systematic bias meanwhile maintaining underlying biological heterogeneity.

**Methods:** To robustly correct the systematic bias, we used the density-power-weight method to down-weigh outliers and extended the one-dimensional robust fitting method described in the previous work of (Windham, 1995, Fujisawa and Eguchi, 2008) to our structured data. We then constructed a robustness criterion and developed a new normalization algorithm, called RobNorm.

**Results:** In simulation studies and analysis of real data from the genotype-tissue expression (GTEx) project, we compared and evaluated the performance of RobNorm against other normalization methods. We found that the RobNorm approach exhibits the greatest reduction in systematic bias while maintaining across-tissue variation, especially for datasets from highly heterogeneous samples.

**Availability:** https://github.com/mwgrassgreen/RobNorm

## 1 Introduction

Mass spectrometry (MS) has made a significant progress over the last few decades, enabling the identification and quantification of thousands or ten thousands of proteins (Callister, et al., 2006; Chawade, et al., 2014; Välikangas, et al., 2018). Nowadays, the use of mass tags allows for multiplexing several samples in a single MS experiment, which permits quantification of protein levels and increases data throughput. This simultaneous measurement also benefits statistical analysis in reducing within-run technical variation. Despite the advancement of the underlying technology and labeled experiment designs, MS data is still affected by the systematic biases introduced during sample preparation and data generation processes (Chawade, et al., 2014). Inclusion of a normalization step is thus needed to correct such systematic biases and to make sample expression more comparable.

Using tandem mass tag (TMT) liquid chromatography-mass spectrometry (LC-MS), (Jiang, et al., 2020) quantified 12,627 proteins from 32 normal human tissue types in the genotype-tissue expression project (GTEx). The dynamic range of protein expression profiles between heterogenous tissue types can be quite different from each other. This makes it hard to distinguish between technical variation and biological variation. In this setting, how to correct for inevitable technical variations while maintaining important biological variation becomes challenging at the normalization step.

One approach from experimental design is to resort to spike-ins or house-keeping peptides/proteins controls. However, distinct from genomic analysis methods, there are no well-defined housekeeping proteins across tissues in the proteomics analysis that could be used in a similar way. From the computational perspective, current widely-used normalization methods for MS data analysis are primarily derived from microarray analysis (Callister, et al., 2006; Chawade, et al., 2014; Välikangas, et al., 2018). Most of these methods rely on the implicit assumption that the protein distributions across all samples are similar. However, this assumption does not hold when expression profiles across or within samples are highly heterogenous, such as from various tissue types in the GTEx project. This motivated us to develop a new data-driven robust normalization method, called **RobNorm**, to robustly correct technical variations while preserving important heterogeneity information.

We used the density-power-weight method to down-weigh outliers and extended the previous work of (Windham, 1995, Fujisawa and Eguchi, 2008) to our setting in **Section 2**. We compared the performance of RobNorm with several commonly used normalization methods in **Section 3**. Not all the normalization methods have the ability to robustly correct the systematic bias without ruining the underlying heterogeneities. Our RobNorm approach showed the best performance for preserving heterogenous sample expression, as shown in **Section 3**. We conclude the paper and discuss a few limitations of the method in **Section 4**.

In this work, we focused on normalizing the relative abundance (in the logarithm scale) of the labeled quantitative proteomics data. Potential application in the label-free quantitative proteomics is discussed at the end of the paper.

## 2 Methods

In our approach, we called the systematic bias the “sample effect”. The protein expression matrix was viewed as structured data with row expression determined by the effect from each protein’s level while column expression was determined by the sample effect. Besides sample effect, protein expression was modeled from a mixture distribution with a Gaussian population distribution. Using this mixture model, we extended the one-dimensional robust fitting shown in the previous work of (Windham, 1995) and (Fujisawa and Eguchi, 2008) to the structured proteomics data and obtained a robust estimation for the sample effect from our algorithm RobNorm. The technical details are provided in the following subsections. In notation below, variables in bold represent vectors and variables with a capital letter denote a matrix based on the context.

### 2.1 Mixture model

Define the protein expression matrix by *X* in log scale. The element *X*_*ij*_ represents the expression for *i*^*th*^ protein from *j*^*th*^ sample, *i* = 1, …, *n*, *j* = 1, …, *m*, where *n* is the protein number and *m* is the sample size. For the *i*^*th*^ protein, the expression *X*_*ij*_ from sample *j* is affected by the sample effect *v*_*j*_. The sample effect *v*_*j*_ is the common factor in all expression data from sample *j*, which is the systematic bias to remove. Besides the sample effect, the assumption that all the sample expression come from the same distribution is hard to uphold using heterogenous samples. Hence, we modeled protein expression from a mixture distribution, allowing the presence of outliers. For each protein, the majority of its expression is determined by the same population distribution. We took a parametric approach to model the overall population distribution as a Gaussian distribution. The remaining expression data are outliers, which can be technical errors or tissue specific expression. Usually their distribution is unknown. Therefore, we did not specify the outlier distribution. This is one advantage of our model. In formula (1), the Gaussian-population mixture model for the expression in the *i*^*th*^ protein is as follows,

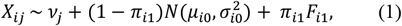

where *π*_*i*1_ ∈ [0, 0.5) is the outlier proportion. From Model (1), a fraction (1 − *π*_*i*1_) of samples come from the Gaussian-population distribution 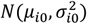 with mean *μ*_*i*0_ and variance 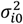, while the rest of samples are outliers from unknown distribution *F*_*i*1_. The parameters 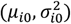 in the Gaussian population are called the protein effect. They can be different in various proteins. If a protein has no outliers, that is *π*_*i*1_ = 0, then its expression can be written as

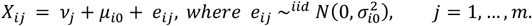

Relaxing this assumption on the outlier distribution makes the model more flexible. Here we assumed independence for all the expression data.

### 2.2 Robust criterion for the structured data

In the presence of outliers, our goal is to robustly estimate the sample effect. The literature of robust estimation is rich in statistics (Hampel, et al., 2011; Huber, 2011; Maronna, et al., 2018; Tyler, 2008). The work of (Basu, et al., 1998; Fujisawa and Eguchi, 2008; Windham, 1995) used an approach to down-weight the outliers by weighting each data point by the fitted density to power *γ*. The parameter *γ* is the exponent of the weighted density. Previous work found that this approach can still maintain robustness even when the outlier proportion is not small, which fits our setting with sample expression that is highly heterogeneous.

Suppose the sample effect *v*_._ is known and then the adjusted expression 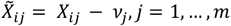, are independently and identically distributed (i.i.d.) from the mixture distribution 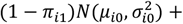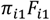. In the procedure of (Windham, 1995), a weight *w*_*ij*_ is assigned to the adjusted expression 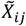 by

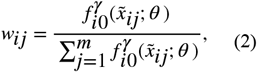

where *γ* ≥ 0, and 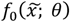 is a Gaussian density function with parameter *θ* = (*μ*, *σ*^2^). If the underlying population parameter is known, that is, 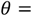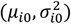, then the theoretical distribution of the weighted data 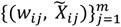, is still Gaussian 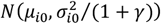, but its variance is shrunk by 1/(1 + *γ*) of the original variance. In the illustrated example shown in **Figure 1**, the outliers go to the tail of the density of the weighted data and thus do not contribute substantially to the population estimation. In this way, the expression from the population gains more weights while the outliers gain less, which achieves the goal of robustness.

**Figure 1:**
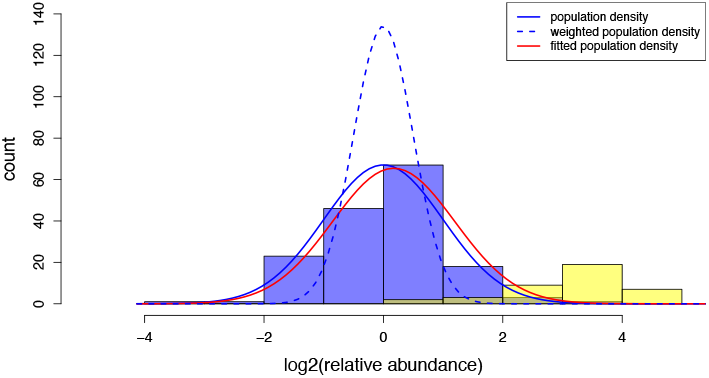
An illustration of the population estimation from the down-weighting outlier procedure using simulated expression for one protein. The protein expression profile is generated from the mixture model 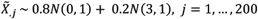. The blue bars (in the left bump) correspond to the population expression (about 80% samples) from distribution N(0, 1) and the yellow ones (in the right bump) correspond to the outlier expression (the rest about 20% from distribution N(3, 1). The blue solid curve indicates the underlying population Gaussian distribution N(0, 1). The blue dashed curve indicates the theoretical weighted population Gaussian distribution N(0, 1/(1 + γ)) from the down-weighting outlier procedure. The red curve is the fitted population distribution from method RobNorm under γ = 1.

Windham’s procedure estimates the population parameters by solving the estimation equation,

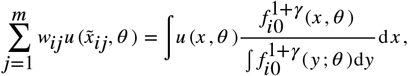

where *u*(*x*, *θ*) = ∂ log *f*_*i*0_(*x*; *θ*)/∂ *θ* is the score function of the loglikelihood function. In the same approach of down-weighting the outliers, the work of (Fujisawa and Eguchi, 2008) found a robustness criterion --- *γ*-cross entropy in (3), which gives the same estimates as from the Windham’s procedure,

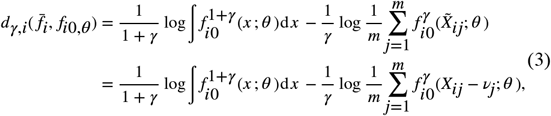

where *γ* > 0 and 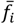 is the empirical density of the adjusted expression in the *i*^*th*^ protein. As *γ* approaches to zero, the limit of the *γ*-cross entropy criterion is the negative of averaged log-likelihood function,

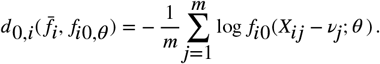

In the case of *γ* = 0, the weight in (2) is 1/*m* for all samples. If there are no outliers, taking *γ* = 0 gives the most efficient estimates --- maximum likelihood estimation (MLE). In the presence of outliers, large *γ* values down-weigh the outliers more aggressively and hence could provide more robustness. The model parameter *γ* essentially balances robustness and efficiency.

To estimate the sample effect in our structured data, we extended the criterion of *γ*-cross entropy for a single protein to the weighted summation of the *γ*-cross entropies from all proteins. Note that *W*_*ij*_ values are self-standardized for each protein, i.e., 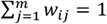. We define the **weighted sample size** by

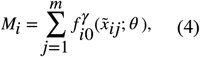

from the denominator of *W*_*i*_ defined in (2). For example, there are five data points {*x*_1_, *x*_2_, …, *x*_5_}. If we re-weight *x*_1_ and *x*_2_ each by 1/2 and others by 0, then the weighted sample size *M* = 2. Our robustness criterion for the structured data is

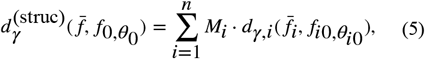

where *d*_γ,i_ is defined in (3) and *M*_i_ is defined in (4). When *γ* = 0, the criterion (5) becomes the negative of log-likelihood function from all the expression.

### 2.3 Robust normalization

Based on criterion (5), the robust estimate for (***v***, ***θ***_0_) is

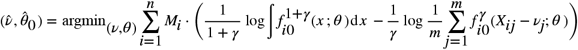

where 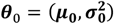 are the population parameters. The weight and the estimates are obtained in an iterative fashion. Given the weights ***W***, taking the derivatives of 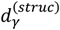 with respect to the parameters gives

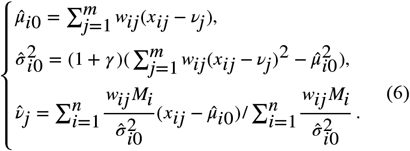

In turn, the weight ***w*** is updated based on 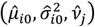 from the previous iteration. The fixed points from the iterations are the final estimates. We summarized the steps in **Algorithm 1**. Note that there is an unidentifiability in estimating *μ*_*i*0_ and *v*_*j*_. Both (*μ*_*i*0_, *v*_*j*_) and (*μ*_*i*0_ − *c*, *v*_*j*_ + *c*) satisfy the equations in (6), where *c* is a constant. One common way to remove this ambiguity is to set *c* as the sample effect for the standard sample ***x***_0_, which is constructed of sample medians from individual proteins. Although our estimation does not rely on such a standard sample, we introduced ***x***_0_ in the algorithm. In the initial step, the estimate for the sample effect ***v***^(0)^ is attained from a commonly used method --- probabilistic quotient normalization (PQN) (Dieterle, et al., 2006). More analysis and performance comparisons with PQN are in the **Supplementary Material**.

**Algorithm 1:**
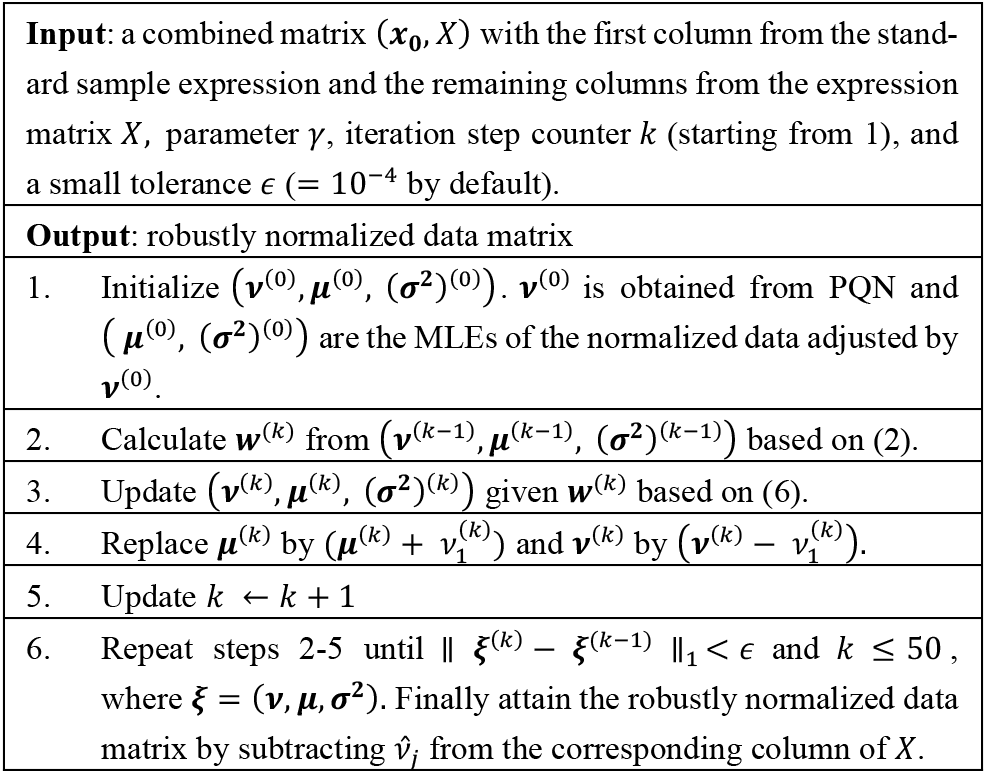
Robust normalization (RobNorm)

## 3 Results

In this section, we first reviewed several existing normalization methods and then compared their performance against our RobNorm approach in both simulation studies and application to real data.

### 3.1 Summary of current normalization methods

Most normalization methods used for the MS proteomics data are adapted from microarray analysis. The work of (Callister, et al., 2006; Chawade, et al., 2014; Välikangas, et al., 2018) gave a systematic review on commonly used normalization methods. Here we summarized these methods into four categories in supplementary **Table S1**.

From TableS1, Category I includes normalization methods that use simple sample shifting, including mean/median normalization and PQN. The advantage of this approach is easy to implement. However, without a targeted model, it is hard to reduce the systematic bias to the desired level.

Our RobNorm method belongs to the category II: model-based normalization. This category also contains the ANOVA-based normalization methods (Hill, et al., 2008; Oberg and Mahoney, 2012; Oberg, et al., 2008) and EigenMS (Karpievitch, et al., 2009). ANOVA-based normalization methods attempt to remove all sources of biases from the model, such as the effects from tags, experimental groups, and peptides. However, aiming to include all these effects in the model may lead to the problem of over-fitting. Moreover, it is impossible to identify all the relevant sources of biases due to the complexity of MS experiments (Karpievitch, et al., 2009). Furthermore, if there is some effect, such as the tissue effect, not included in the model, the estimation of the sample effect from ANOVA may not be robust. EigenMS is another approach adapting the surrogate variable analysis (SVA) method of (Leek and Storey, 2007) to remove possible bias from unmodeled effects. It was formulated in two steps: (i) to remove the known effects from experimental design, and (ii) to apply SVA on the residuals to remove possible unknown bias trends. One concern inherent to this method is that it may eliminate the biological differences in the differentially expressed (DE) proteins, especially when the truly differential expression is strong and dense. Since EigenMS was implemented in two successive steps, the robustness of its estimation in the first step may affect the stability of its results in the second step. Compared to ANOVA-based normalization that includes all effects in one model and EigenMS that estimates the known and unknown effects in two steps, we instead put unmodeled and unknown factors into the outlier distribution and then robustly estimated the sample effect.

Category III comprises the sample variance stabilization normalization (VSN) method (Huber, et al., 2002). Methods based on sample-to-reference transformation belong to Category IV. The assumption of the methods in these two categories is that the majority of the proteins are non-differentially expressed (nonDE). Since nonDE proteins are mainly affected by systematic bias, the normalization methods attempt to correct this bias based on nonDE protein data. As the nonDE proteins are unknown beforehand, several normalization methods make use of robust estimation approaches. Quantile normalization is based on this assumption and transforms the distributions of all samples to be conform to the empirical cumulative density function (e.c.d.f.) of the reference sample (Bolstad, et al., 2003). VSN estimates the sample transformation parameters from the least trimmed sum of squares (LTS) regression. In its implementation, VSN provides options for tuning the LTS quantile parameter for robustness. Linear regression-based normalization methods use robust least median regression (rlm) to project the unnormalized expression onto the reference sample expression (Chawade, et al., 2014). Loess-based methods are based on robust loess fitting tuned by a span parameter (Ballman, et al., 2004; Dudoit, et al., 2002; Ting, et al., 2009). However, when the sample expression is highly heterogeneous, the assumption that the majority of the proteins are nonDE is hard to maintain. In the simulation studies in **Section 3.2**, we investigated the performance of these robustness methods against differing degrees of heterogeneity to identify the level of heterogeneity at which methods become unreliable.

### 3.2 Simulation studies

We examined the performance of the normalization methods mentioned in **Section 3.1** in simulated data. We listed the methods for comparison below and provided their implementations in R functions (Team, 2013) as references. The methods for comparisons are: our RobNorm approach under *γ* = 0.5,1 for the large sample size and under *γ* = 0.1,0.5 for small sample size, ANOVA method (*γ* = 0), mean/median normalization, PQN (Dieterle, et al., 2006), EigenMS (Karpievitch, et al., 2009), VSN (R library vsn – justvsn(.) (Huber, et al., 2002)) under LTS quantile parameter 0.9, 0.7, 0.5, quantile normalization (R library preprocessCore – normalize.quantiles(.) (Bolstad, et al., 2003)), Rlr and its variant RlrMA (Chawade, et al., 2014), LoessCyc and its variant LoessMA under span parameters 0.9, 0.7, 0.5 (R library limma – normalizyCyclicLoess(.) (Ritchie, et al., 2015)). For the Loess-based normalization (Dudoit, et al., 2002), we used fast Loess for computational efficiency.

#### Simulated data

Using the same notation as in model (1), the outlier distribution *F*_*i*1_ here is specified as Gaussian 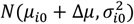. The generative underlying model in the simulations is as follows

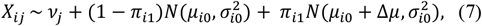

where *i* = 1, …, *n*, and *j* = 1, …, *m*. The protein population mean *μ*_*i*0_’s were generated from *N*(0,1) and the protein variance 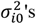 were from inverse-Gamma distribution with the shape parameter at 5 and scale parameter at 0.5. 80% of sample effect *v*_*j*_’s were obtained from *N*(0,1), and the remaining 20% *v*_*j*_’s from *N*(1,1). The distributions of the generated parameters are shown in supplemental Figure S1. The first half of the samples were treated in group 1 and the last half in group 2. In the simulations, the outliers were concentrated in two regulation blocks, one block per group. The simulated dataset was visualized in **Figure 2** (left panel). The 80% expression levels in the up-regulated block (in the upper left corner in color red) was up-shifted by Δ*μ* in mean and, similarly, 80% expression levels in the down-regulated block (in the bottom right corner in color blue) were down-shifted by Δ*μ* in mean. There is no overlap between the regulation blocks.

**Figure 2:**
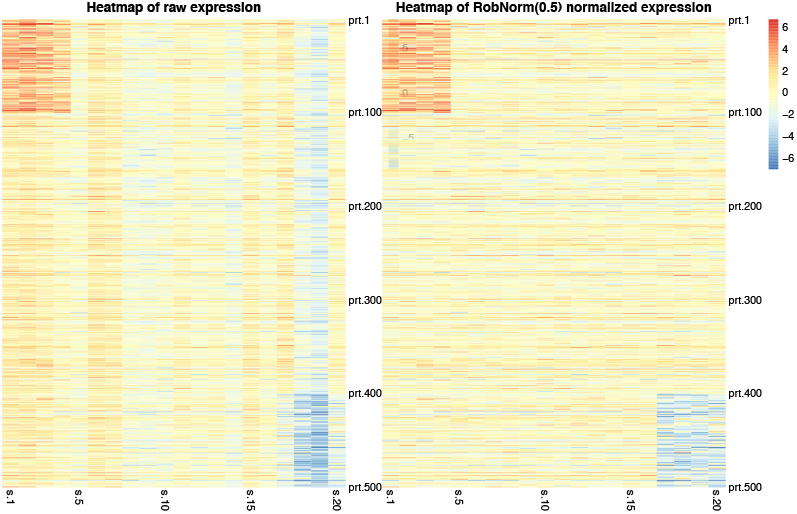
An illustration of a simulated data expression matrix under the setting of protein number n = 500, the sample number m = 20 and the regulation effect |Δμ| = 3. Each row is the expression of a protein from 20 samples and each column is the expression of a sample from 500 proteins. There are two regulation blocks. One occurs in the upper left block in the first 100 proteins from the first four samples and the other in the bottom right block from the last 100 proteins from the last four samples. Each sample (the column) is affected by a sample effect. The left panel is the raw expression. The right panel is the normalized expression from method RobNorm under parameter γ = 0.5. The simulation details are in Section 3.1.

In the simulated data shown in **Figure 2**, there are clear stripes in the columns, indicative of the sample effect. The right panel in **Figure 2** shows the normalized expression from RobNorm. After the sample effect was robustly removed, the underlying regulation blocks were recovered.

In the differential expression (DE) analysis, four cases were considered: (i) DE protein proportion = 2×10% (10% for up-regulation, 10% for downregulation) and DE mean change Δ*μ* = ±1, (ii) DE protein proportion = 2×10% and Δ*μ* = ±3, (iii) DE protein proportion = 2×20% and Δ*μ* = ±1, (iv) DE protein proportion = 2×20% and Δ*μ* = ±3. The protein size *n* was set as 5000 and the sample size *m* as 200 and 40. The Wilcoxon rank sum test was applied for each protein after normalization and the Area Under the Curve (AUC) for each method was recorded. We repeated the procedure independently 20 times. The reports were summarized in **Figure 3** (*m* = 200) and supplementary Figure S2 (*m* = 40).

**Figure 3:**
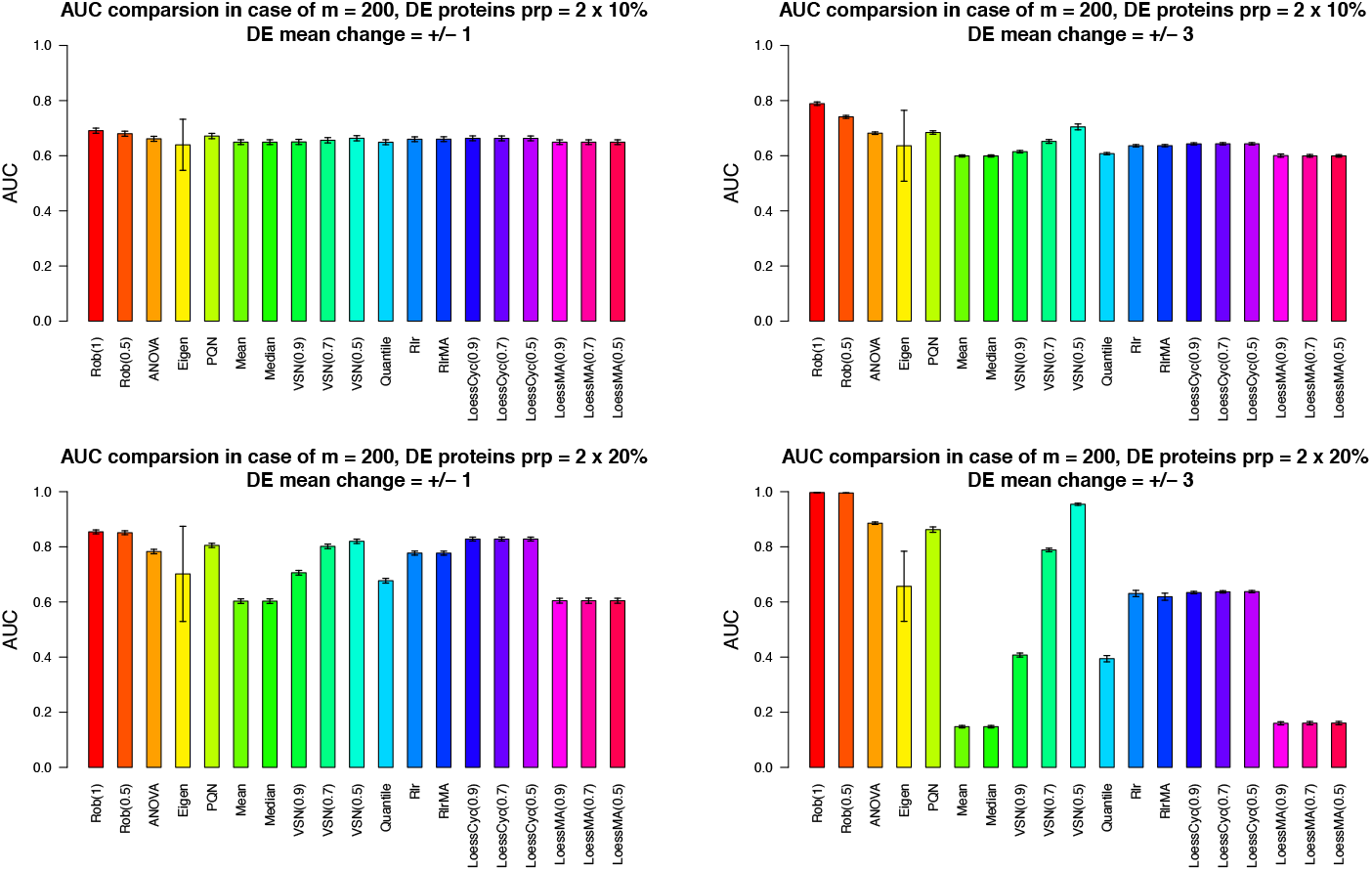
The effect of normalization method on the differential expression (DE) analysis in terms of AUC from simulation studies. In the simulations, the sample size m was set as 200. Four situations were considered and each panel shows the AUC results in each situation. Situation (1):DE protein proportion = 2 × 10% and DE mean change = ±1 (the topleft panel); Situation (2): DE protein proportion = 2 × 10% and DE mean change = ±3 (the topright panel); Situation (3): DE protein proportion = 2 × 20% and DE mean change = ±1 (the bottomleft panel); Situation (4): DE protein proportion = 2 × 20% and DE mean change = ±3 (the bottomright panel). The height of each bar shows the average AUCs from 20 independently repeated simulations. The error bar corresponds to ±1 standard deviation from the averaged AUC.

When sample size *m* = 200, RobNorm delivered the best performance in all four cases (**Figure 3)**. The performance of RobNorm under *γ* = 1 was slightly better than that under *γ* = 0.5, especially in the case that the outlier magnitude was large. Among the methods in Category I, PQN performed the best but was still out-performed by RobNorm, which indicated the importance of incorporating data structure into the normalization step. Using quantile parameters from 0.9 to 0.5, the performance of VSN was dramatically improved. Under quantile parameter = 0.5, VSN removed 50% extreme data points to be robust enough to estimate the normalization factors, although it still performed worse than RobNorm. Other methods in Category II generally performed worse than RobNorm. Without iterating the expression of the reference sample, the LoessMA had even lower power than LoessCyc at finding DE proteins. One observation in this simulation study was that the results from EigenMS had much larger variation than all other approaches. We speculated that this variation may arise from failures of distinguishing true signals from unwanted bias.

Comparison results under sample size *m* = 40 are summarized in **Supplemental Material**. When the sample size was small, the fitted population of some proteins could be locally trapped such that the variance of those proteins was very small under a large *γ*. To avoid this, a small *γ* for RobNorm is recommended. In simulations where *m* = 40, we set *γ* = 0.1, 0.5. RobNorm under *γ* = 0.5 performed better than the rest of the methods except EigenMS. EigenMS achieved the best performance on average but at the cost of larger variation. Note that EigenMS adjusted both known and unknown biases while other methods focused only on adjusting known biases.

We further investigated the performance of RobNorm and one competitive method PQN in estimating the sample effect and protein effect under various outlier proportions and magnitudes. The effect of the choice of *γ* was also explored in **Supplementary Material.** In the simulation studies under *m* = 200, the estimation for the sample effect from RobNorm achieved high accuracy in terms of sum of squared errors and was not affected much by the choice of *γ*. We observed that a properly selected *γ* would improve the estimation accuracy for the protein effect.

### 3.3 Real data application

We applied the normalization methods above to the labeled proteomics dataset generated in our previous work (Jiang, et al., 2020). In this dataset, each run had a pooled reference sample. The peptide abundances were first normalized by the total sum normalization and then summarized to the protein level. Their relative abundances were obtained by calculating the ratios of the sample abundances to the reference. To evaluate normalization performance, we only considered the 5,970 proteins that were observed in at least 100 samples to avoid possible bias from missing values. Details in identification and quantification can be found in the Supplementary Material in (Jiang, et al., 2020).

With the exception of the VSN method working with the raw relative abundance data, all the other normalization methods use log transformed data. For the fast loess (LoessCyc) normalization method, the default iteration limit is 3. We found that the estimation from LoessCyc did not converge within 100 iterations for this dataset. Hence, we did not include LoessCyc and only included the LoessMA method which does not iterate reference sample expression. The methods applied in this dataset were RobNorm (under *γ* = 0.5, 1), ANOVA (*γ* = 0), EigenMS, PQN, mean/median normalization, quantile normalization, Rlr, RlrMA, VSN (under quantile parameter = 0.9,0.7,0.5), and LoessMA (under span parameter = 0.9,0.7,0.5). The implementation of EigenMS removed the proteins with any missing values so the normalized data from EigenMS covered only 4,816 proteins.

To evaluate the performance of these normalization methods, (Callister, et al., 2006; Chawade, et al., 2014; Välikangas, et al., 2018) discussed several relevant metrics, such as quantitative metrics to evaluate within-group and across-group variation and qualitative visualization measures. We first evaluated the effect of the normalization method on the within-tissue variation in terms of the pooled intragroup median absolute deviation (PMAD). From **Figure 4**, except the EigenMS method, most of the methods including RobNorm did not significantly decrease the PMADs compared to the raw unnormalized data. It may be because the high quality of the reference samples and labeled experimental design help reduce within-tissue variation in the raw data. We next focused on evaluating the effect of the normalization method on the across-tissue variation.

**Figure 4:**
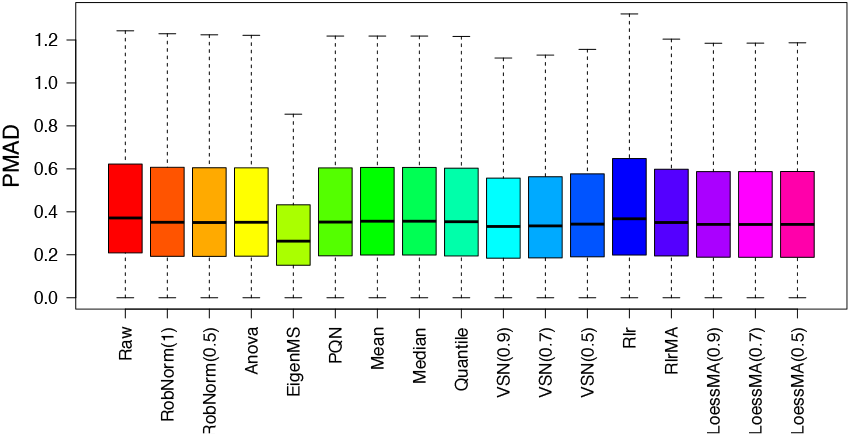
The effect of normalization method on within-tissue variation from the GTEx proteomics dataset. The within-tissue variation was measured by the pooled intragroup median absolute deviation (PMAD) method introduced in (Chawade, et al., 2014).

As a qualitative measure, we visualized the distribution of tissue expression after normalization. In **Figure 5**, the densities of muscle and heart ventricle expression obviously lagged behind other tissue expression towards low values in the raw data. This can be explained by the fact that high signals from a small number of abundant proteins can suppress the signals from lower abundance proteins in muscle and heart tissues (Geiger, et al., 2013; Jiang, et al., 2020; Wang, et al., 2019). The purpose of normalization is to make most of the expression comparable across all tissues and the adjustment factors should not be strongly affected by a few extremely high or low abundances. In RobNorm normalized data, the density peaks of all the tissues are aligned together in **Figure 5,** while the EigenMS and VSN adjusted data still produce muscle and heart tissue densities deviated from other tissue densities, which lowers their ability to detect up-regulated muscle or heart proteins. All method density comparisons are shown in supplementary Figure S9.

**Figure 5:**
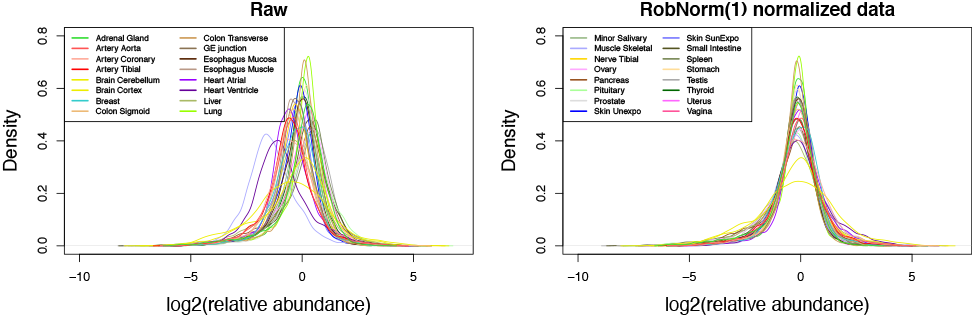
The distribution of relative protein abundances in log scale at base 2 across tissues. The protein expression from each tissue is summarized from sample medians in the same tissue. The left panel shows the tissue densities from the raw dataset before normalization. The right panel shows the densities from the normalized dataset from method RobNorm under γ = 1. The density plots from all the normalized methods are shown in Figure S9. The legends for the tissue types are divided in the two panels.

We further investigated the normalization effect on the across-tissue variation in differential expression (DE) analysis. As full tissue expression analysis was studied in (Jiang, et al., 2020), here we focused on comparing muscle group expression vs. non-muscle group expression on the normalized datasets. The proteins observed in at least five samples in both two groups were included in this DE analysis. The Wilcoxon rank sum test was applied for each protein and significantly regulated proteins under BH adjusted p-value < 0.05 were reported (Benjamini and Hochberg, 1995). The DE results are summarized in the supplementary Figure S10 with volcano plots. Based on the significantly up-regulated muscle proteins from the DE results, GO term biological function analysis was applied and was implemented using the software STRING (Franceschini, et al., 2012(Franceschini, et al., 2012)). 1,232 significantly up-regulated proteins resolved in muscle were detected from the RobNorm adjusted dataset. They were significantly enriched in the skeletal-muscle-function-related GO terms, including respiratory electron transport chain, oxidative phosphorylation, mitochondrial electron transport, NADH to ubiquinone, muscle system processes, muscle contraction, aerobic respiration, glycolysis, and fatty acid metabolic process (with BH adjusted p-value < 10^−8^). Approximately 900 proteins detected using RobNorm were not detected in raw data or in EigenMS adjusted data. Those undetected proteins were highly enriched in muscle-related functions, including generation of precursor metabolites and energy, muscle system process, muscle organ development, muscle contraction, mitochondrial electron transport, NADH to ubiquinone, and skeletal muscle contraction (with BH adjusted p-value < 10^−5^). This indicated that improper normalization methods may limit discoveries in protein function and tissue regulation. Moreover, we evaluated the coverage of significantly up-regulated proteins over the proteins belonging to four well-known muscle-function related GO terms -- NADH ubiquinone oxidoreductase subunit, myosin, mitochondrial protein, and ATP protein. As shown in **Figure 6**, the coverage from raw data, EigenMS, VSN, Quantile, mean, and median normalized data are less than 80% in at least one protein group. The Rlr method and our RobNorm approach have similar coverage. For the down-regulated proteins, we found that those proteins whether they were detected from RobNorm or other methods were mainly enriched in the basic protein functions, such as vesicle-mediated transport, protein transport, and protein localization. Moreover, we did not obtain much significant GO term enrichment associated with non-muscle-tissue-type specific functions. It may be because we restricted the proteins to be observed in at least five samples in each group in this DE analysis such that the non-muscle-tissue-type specific proteins were filtered out. Hence, here we did not further evaluate performance differences in detecting down-regulated proteins.

**Figure 6:**
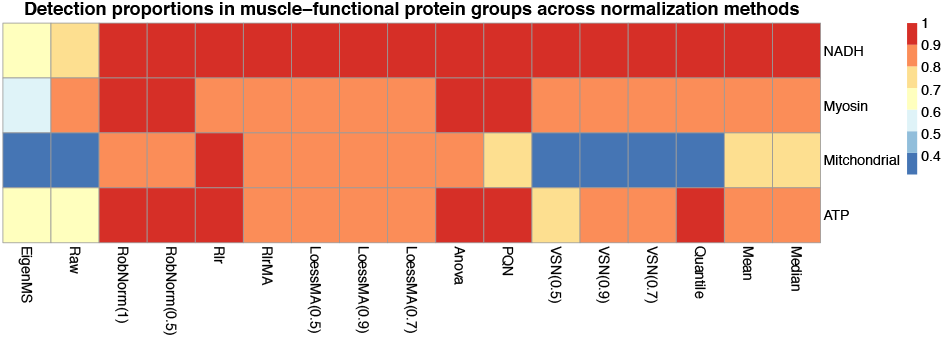
The effect of normalization method on muscle-function related GO terms using only the up-regulated proteins for each method from differential expression (DE) analysis. The muscle-function related four GO terms were pre-selected, including NADH ubiquinone oxidoreductase subunit, myosin, mitochondrial protein, and ATP protein. Each cell in the heatmap shows the proportion of the number of significant DE muscle proteins within one GO term over the total number of the proteins in the corresponding GO term. The DE muscle proteins were obtained from Wilcoxon rank sum test for muscle sample expression versus non-muscle sample expression comparison on the raw or normalized dataset.

## 4 Discussion and conclusion

In analyzing MS data from labeled experimental designs, we developed a new data-driven robust normalization method (RobNorm) and compared the performance of RobNorm to several commonly used normalization methods. From these studies and their application to real data, we concluded that our RobNorm approach offers the best performance in correcting systematic bias while maintaining underlying biological heterogeneities. However, there are still some limitations in the method and future work to be done.

### Model assumption

The RobNorm approach was based on the assumption that the majority of protein expression follows a Gaussian distribution in logarithmic scale. Based on the Gaussian population assumption, we obtained an explicit formula to estimate the sample effect. If the underlying distribution has heavier tails, such as a *t*-distribution with a small degree of freedom, a similar framework can be still applied. To maintain the power for performance at normalization, it is required to adjust the weight function based on the *t*-distribution.

### Model flexibility and stability

To model the protein expression in the step of normalization, we only modeled the sample effect and the protein effect as the primary parameters, while EigenMS considered both known and unknown effects. As pointed out in (Karpievitch, et al., 2009), normalization models need to be flexible enough to capture biases of arbitrary complexity while avoiding overfitting that would invalidate downstream statistical inference. From our simulation studies with highly heterogenous sample expression, we found that EigenMS failed to distinguish the true signals from the unwanted effects and its performance had high variation. There is still a need to robustly and stably remove both known and unknown systematic biases in future work.

### Choice of γ

Our robust estimation was based on the density-power-weight approach. The model parameter *γ* is the weight exponent, which balances robustness and efficiency of the estimation. Since our algorithm sets the same *γ* for all proteins, the *γ* can be large for some proteins such that their population fitting was locally trapped, i.e., the estimated variance was too small. To avoid this, we suggested choosing a smaller *γ* when the sample size is small. Therefore, in implementing RobNorm from GitHub, we included a warning message if the prechosen γ is large. From our experience, we recommended setting *γ* as 1 or 0.5 when sample size is greater than 100, and otherwise setting *γ* as 0.5 or 0.1. The ideal case is to choose *γ* adaptively for each protein, but this will lead to a problem in balancing flexibility and stability of the model. How to select an optimal *γ* is still an open and interesting problem.

### Sample size

Since our robust estimation for the sample effect depends on the estimation of the population parameters, the sample size cannot be too small. This is one limitation of the RobNorm method. In practice, we suggest that the sample size should be greater than or equal to 20.

### Missing values

In practice, missing values are very common in MS data. Since RobNorm is based primarily on population expression, random missing values would not have much effect on the normalization factors. If there are missing values in the population, one can impute the missing values by taking the sample median or the robustly fitted mean and then iteratively applying our algorithm until the estimated parameters converge. To avoid possible bias from missing values due to low expression, we recommended using partial proteins with missing proportion < 50% to estimate the sample effect and then apply the estimated sample effect to normalize all the proteins. The work of (Karpievitch, et al., 2012) combined their EigenMS normalization method with missing value imputation. How to embed the missing value imputation step into our framework can be further explored.

### Extension to label-free experimental designs

Our RobNorm method was designed to normalize labeled proteomics data. Label-free proteomics quantification is usually considered to be more noisy by nature when compared to labeled data (Callister, et al., 2006; Cox, et al., 2014). One step normalization may not be enough to correct all the biases. The work of (Kultima, et al., 2009) combined the normalization step with removing run order bias. MaxLFQ took pair-wise comparison of peptides to best estimate protein abundance (Cox, et al., 2014). Also, different search engines may affect the quantification results (Kuharev, et al., 2015). It is possible to apply RobNorm to label-free quantification as long as the Gaussian assumption is valid for the population expression. However, there still needs a combination of multiple processing steps used, not only one normalization step, to fully correct systematic biases.

## Acknowledgements

The authors thank the funding supports by NIH eGTEx grant U01HG007611 (M.P.S., H.T.), NIGMS grant GM073059 (H.T.), NIGMS grant R35GM127063 (H.T.), NIH CEGS grant 2RM1HG00773506, NIH GTEx grant 5U01HL13104203. The authors thank Pier Luigi Martelli, Associate Editor and three reviewers for the insightful comments that greatly improved the article.

## Data availability

The proteomics data used in normalization method comparisons can be found in paper (Jiang, et al., 2020). The code for RobNorm is in GitHub https://github.com/mwgrassgreen/RobNorm.

## Authors’ contributions

MW and HT developed the method. LJ, RJ and JC generated the proteomics data. HT and MPS helped analysing the data. MW wrote the paper and MPS, LJ, QL helped to write. All the authors contributed to discussion and revised the paper.

