## Supplemental Material for "RobNorm: Model-Based Robust Normalization Method for Labeled Quantitative Mass Spectrometry Proteomics Data"

### Supplemental Materials

#### Normalization method summary.

| **Table S1: Normalization Method Summary for Labelled Proteomics Quantification** | |
|---|---|
| **Notation:** | |
| $x_{.j}$: the unnormalized expression<br>in log scale for sample $j$<br>$\mu_i$: main effect for protein $i$ | $\nu_j$: sample effect for sample $j$<br>$\widetilde{x_{.j}}$: normalized expression<br>$\hat{\upsilon}_j$: the adjustment factor |
| **Category I: simple sample shift normalization**<br>Adjustment $\widetilde{x_{.j}} = x_{.j} - \hat{\upsilon}_j$, setting $\hat{\upsilon}_{ref} = 0$ for reference sample $x_{.ref}$<br>$x_{.ref}$ is obtained from protein medians | |
| Mean normalization | $\hat{\upsilon}_j = mean\, x_{.j} - mean\, x_{.ref}$ |
| Median normalization | $\hat{\upsilon}_j = median\, x_{.j} - median\, x_{.ref}$ |
| Probabilistic quotient normalization(PQN) (Dieterle, et al., 2006) | $\hat{\upsilon}_j = median\,(x_{.j} - x_{.ref})$ |
| **Category II: model-based sample normalization**<br>Model $X_{ij} = \nu_j + \mu_i + (other\ effects) + \epsilon_{ij}$ where $\epsilon_{ij}$ is the noise<br>Adjustment $\widetilde{x_{.j}} = x_{.j} - \hat{\upsilon}_j$, | |
| ANOVA normalization (Hill, et al., 2008; Oberg and Mahoney, 2012; Oberg, et al., 2008) | Model $X_{ij} = \nu_j + \mu_i + \xi_{ij} + \epsilon_{ij}$and noise $\epsilon_{ij} \sim^{iid} N(0, \sigma_i^2)$<br>The effect $\xi_{ij}$ may come from experimental effects, peptide effect etc.<br>To estimate $\nu_j$ from least squares |
| EigenMS<br>(Karpievitch, et al., 2009) | Model<br>$X_{ij} = \nu_j + \mu_i + (unknown\ factors) + \epsilon_{ij}$, and noise $\epsilon_{ij} \sim^{iid} N(0, \sigma_i^2)$<br>Adapting SVA to remove unknown factors |
| Our RobNorm | Model $X_{ij} = \nu_j + \mu_i + \epsilon_{ij}$ where<br>$\epsilon_{ij} \sim (1 - \pi_{i1}) N(0, \sigma_{i0}^2) + \pi_{i1} F_{i1}$<br>To estimate $\nu_j$ from robust fitting |
| **Category III: sample variance stabilization normalization**<br>Consider raw expression $y_{ij} = 2^{x_{ij}}$ | |
| VSN<br>(Huber, et al., 2002) | Model $h(y_{ij}) = arcshih(a_j + b_j y_{ij})$<br>$h(y_{ij}) \sim^{iid} N(\mu_i, \sigma^2)$ for nonDE proteins. Adjustment: $\tilde{y}_{ij} = arcshih(\hat{a}_j + \hat{b}_j y_{ij})$ and $(\hat{a}_j, \hat{b}_j)$ estimated from LTS |
| **Category IV: sample-to-reference transformation normalization**<br>Adjustment $\tilde{x}_{.j} = h(x_{.j}, x_{.ref})$<br>where $x_{.ref}$ is obtained from protein medians | |
| Quantile normalization<br>(Bolstad, et al., 2003) | $\tilde{x}_{ij} = F_j^{-1}(Q(x_{i0}))$ where $x_{.0}$ is a baseline sample, Q ($F_j$) is the empirical c.d.f. of $x_{.0}$ $(x_{.j}, resp.)$ |
| Linear regression normalization (Rlr)<br>(Chawade, et al., 2014) | $\tilde{x}_{.j} = (x_{.j} - \hat{a}_j) / \hat{b}_j$, where $(\hat{a}_j, \hat{b}_j)$ estimated from $rlm(x_{.j} \sim a_j + b_j\, x_{.ref})$ |
| Linear regression normalization with MA transformation (RlrMA)<br>(Chawade, et al., 2014) | $\tilde{x}_{.j} = x_{.j} - (\widehat{\Delta x}_{.j})$ where $\Delta x_{.j} = x_{.j} - x_{.ref}\ and\ (\widehat{\Delta x}_{.j})$ estimated from $rlm(\Delta x_{.j} \sim a_j + b_j\, x_{.ref})$ |
| Fast cyclic loess normalization (loessCyc)<br>(Ballman, et al., 2004; Ting, et al., 2009) | $\tilde{x}_{.j} = x_{.j} - (\widehat{\Delta x}_{.j}), (\widehat{\Delta x}_{.j})$ estimated from $loess(\Delta x_{.j} \sim a_j + b_j\, x_{.ref})$ under a certain span parameter.<br>$x_{.ref}^{(t)}$ is updated along iterations. |
| Fast MA loess normalization (loessMA) | The same as loessCyc except without updating $x_{.ref}$ |

#### Simulation studies.

**Robust estimation comparison.** In this study, we focus our comparison on one competitive method PQN. Since both RobNorm and PQN are in a linear correction and they compare to the same standard sample, we compare their estimation accuracy on the same footing. We investigate the performance of our RobNorm and PQN in the four cases defined in the **Section 3.1** in the main text. In the simulatins, **w**e generate simulated data from model (7) in the main text. We set the size of each regulation block as $(20\% \times 5000)$ proteins times $(20\% \times 200)$ samples and the regulation rate in each block as 80%. The standard sample is set as the true protein population mean $\boldsymbol{\mu}_0$. In this way, the estimates $\hat{\boldsymbol{\nu}}$ and $\hat{\boldsymbol{\mu}}_{\mathbf{0}}$ are truly for the underlying $\boldsymbol{\nu}$ and $\boldsymbol{\mu}_{\mathbf{0}}$. We take the Sum of Squared Errors (SSE) to evaluate the accuracy of the estimation. The SSE for $\boldsymbol{\theta}_0$ is defined by $\| \hat{\boldsymbol{\theta}}_{\mathbf{0}} - \boldsymbol{\theta}_0 \|_2^2$, where the $\boldsymbol{\theta}_0$ is for the underlying sample effect and population mean and variance.

We report the estimation results in supplementary Figure S3 – S6 for four cases respectively. We can see that in each case, our robust estimate for the sample effect $\boldsymbol{\nu}$ has lower SSE and shows its advantage especially in the large regulation cases. To estimate $\boldsymbol{\mu}_0$, in the small regulation cases our robust estimates under $\gamma = 0.5, 1$ have similar performances. In the large regulation cases, our robust estimates for $\boldsymbol{\mu}_0$ under $\gamma = 1$ have slightly lower bias than the estimates under a smaller $\gamma = 0.5$. To estimate $\boldsymbol{\sigma}_{\mathbf{0}}^{\mathbf{2}}$, our robust estimate has slightly lower bias under a smaller $\gamma = 0.5$ in the small regulation, while it has much lower bias under a bigger $\gamma = 1$ in the large regulation. This suggests that choosing a proper $\gamma$ is really data dependent. How to choose an optimal $\gamma$ is out of the scope of this paper. Our simulations show that $\gamma = 0.5, 1$ does not have much effect on the accuracy in estimating the sample effects and the protein population means. More estimation comparisons under various sizes and magnitudes of regulated blocks are in supplementary Figure S7 – S8.

**Robust estimation comparison under various sizes and magnitudes of regulation blocks.** We investigate the performances of our RobNorm in various sizes of regulation blocks. For simplicity, here we consider that there is one up-regulated block affecting $(100 \times \text{prt.prp})\% \times 5000$ proteins in $(100 \times \text{s.prp})\% \times 200$ samples and the regulation rate is 80%. We set (prt.prp, s.prp) varying in the grid of $(0.1, 0.2, 0.3, 0.4, 0.5) \times (0.1, 0.2, 0.3, 0.4, 0.5)$. We consider two situations: (i) small regulation with $\Delta\mu = 1$ and (ii) large regulation with $\Delta\mu = 3$. We set $\gamma = 0.5$ for our RobNorm method. We generate the data from the model (7) in the main text under each setting $(\Delta\mu,\ \text{prt.prp},\ \text{s.prt})$. To apply **Algorithm 1**, we take the standard sample as the sample medians from each protein, denoted by $\text{rowMed}(X) =: \boldsymbol{x}_{\mathbf{0}}$. Since $\boldsymbol{x}_{\mathbf{0}}$ could not exactly be the true protein population mean $\boldsymbol{\mu}_{\mathbf{0}}$, what our RobNorm and PQN estimate are $\boldsymbol{\mu}_{\mathbf{0}} + c$ and $\boldsymbol{\nu} - c$, where c is a constant. Hence, to evaluate the performances of the estimators, we consider

$$(\text{error in } \nu) = \frac{1}{m} \sum_{k=1}^{m} \frac{1}{m-1} \|(\hat{\nu} - \hat{\nu}_k) - (\nu - \nu_k)\|_2^2, \quad (1)$$

and

$$(\text{error in } \mu) = \frac{1}{n} \sum_{k=1}^{n} \frac{1}{n-1} \|(\hat{\mu} - \hat{\mu}_k) - (\mu - \mu_k)\|_2^2. \quad (2)$$

For $(\text{error in } \boldsymbol{\nu})$, the term $(\hat{\boldsymbol{\nu}} - \hat{\nu}_k)$ - $(\boldsymbol{\nu} - \hat{\nu}_k)$ eliminates the ambiguous constant. We average the sum of squared errors across all the elements and further take the average of the errors based on different $\nu_k$'s. The same argument applies to $(\text{error in } \boldsymbol{\mu})$, in (2).

We simulate data 20 times and summarize the results in supplemental Figure S7 -- S8. In supplemental Figure S7, we compare the performances of RobNorm to PQN in estimating the sample effect $\boldsymbol{\nu}$. In each panel, the first column $(\text{s.prp} = 0)$ and the last row $(\text{prt.prp} = 0)$ simulate the case of no outliers at all. In each cell of the heatmap, the first line is the average error and the second line is the standard deviation of the estimates from 20 repeats. With increasing the proportions in the affected proteins and samples, the error increases, as expected. From supplemental Figure S8, we can see that in the case of small regulation (the panels in the first row), the performance of RobNorm is a little better than that of PQN, while in the case of large regulation, RobNorm outperforms PQN. Supplemental Figure S7 highlights the regions from RobNorm that have average errors less than $10/n = 0.002$ (in color yellow), which are much larger than those from PQN. From the simulations, our robust normalization method still keeps high accuracy under heavy outliers, especially when the signals of the outliers are strong. Similarly, comparing the estimation accuracy for the protein population means shown supplemental Figure S8, our RobNorm performs comparable to the MLEs after PQN adjustment in the case of small regulation. In the case of large regulation, the estimation from RobNorm has larger high accuracy regions (under the threshold $10/m = 0.05$).

#### Normalization comparison to PQN

Consider sample $\boldsymbol{x}_j$ to be written as $(\nu_j\ \mathbf{1} + \boldsymbol{\mu_0} + \boldsymbol{s_j} + \boldsymbol{e_j})$ where $\mathbf{1}$ is an all-ones vector, $\boldsymbol{s_j}$ is a zero vector except nonzero in some regulated proteins and $\boldsymbol{e_j}$ is the Gaussian noise with mean 0. Define the regulation matrix $S = (\boldsymbol{s_1}, \dots, \boldsymbol{s_m})$ and the noise matrix $E = (\boldsymbol{e}_1, \dots, \boldsymbol{e}_m)$. Suppose the mean of the regulations does not affect the protein medians much. We have

$$\boldsymbol{x}_0 = \boldsymbol{\mu}_0 + \text{rowMed}(\mathbf{1} \cdot \boldsymbol{\nu}^T + S + E) \approx \boldsymbol{\mu}_0 + c\ \mathbf{1} + \boldsymbol{e_c},$$

where $c = Med(\boldsymbol{\nu})$ and $\boldsymbol{e_c}$ is the Gaussian noise. The amount PQN adjusts in sample $\boldsymbol{x}_j$ is

$$\text{Med}(\boldsymbol{x}_j - \boldsymbol{x}_0) \approx \text{Med}\left((\boldsymbol{\mu}_0 + \nu_j\ \mathbf{1} + \boldsymbol{s_j} + \boldsymbol{e_c}) - (\boldsymbol{\mu}_0 + c\ \mathbf{1} + \boldsymbol{e_c})\right) = \text{Med}\left((\nu_j - c)\mathbf{1} + \boldsymbol{s_j} + \boldsymbol{e_c}\right) = \nu_j - c + (\Delta c)_j,$$

where $(\Delta c)_j = Med(\boldsymbol{s_j} + \boldsymbol{e_c})$. Hence, the adjusted sample is

$$\boldsymbol{x}_j^{(adj)} = \boldsymbol{x}_j - \left(\text{Med}(x_j - \boldsymbol{x}_0)\right)\mathbf{1} \approx (\boldsymbol{\mu}_0 + \nu_j\ \mathbf{1} + \boldsymbol{e_j}) - \left(\nu_j - c + (\Delta c)_j\right)\mathbf{1} \approx \boldsymbol{\mu}_0 + \left(c + (\Delta c)_j\right)\mathbf{1} + \boldsymbol{e_j}$$

We can see the term $\Delta c$ varies in samples. When the signal effect is strong and the noise variation is large, what PQN adjusts is no longer the systematical sample effect, but partial of the regulation effects. However, our RobNorm automatically captures the distribution of the majority points while the majority points are not affected by the regulation under our assumption. In this way, our method distinguishes the expressions $(\boldsymbol{\mu}_0 + c\mathbf{1} + \boldsymbol{e}_j)$'s from few regulated expressions $(\boldsymbol{\mu}_0 + \left(c + (\Delta c)_j\right)\mathbf{1} + \boldsymbol{e}_j)$'s. That is why our method can be more robust than PQN in this case.

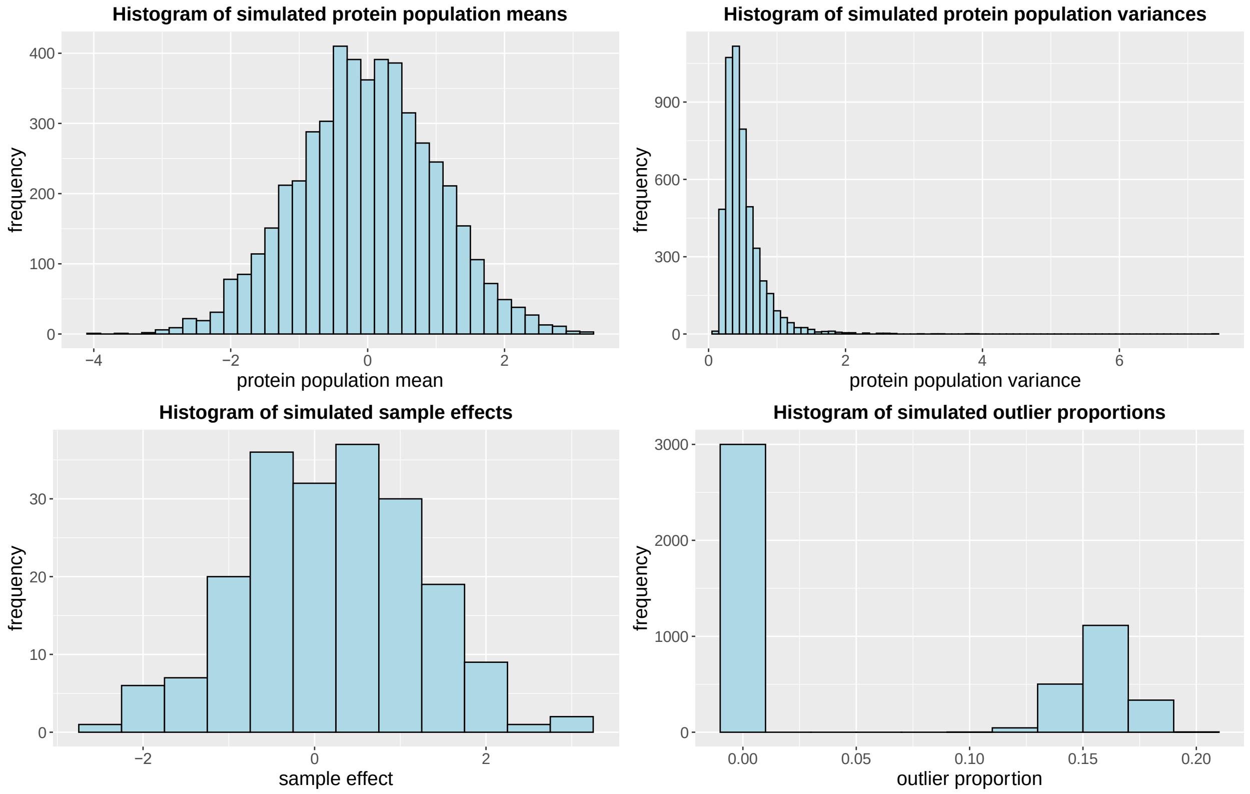


*Figure S1: Histogram of the simulated parameters from model (1) under n=5000, m=200 in one simulation.*

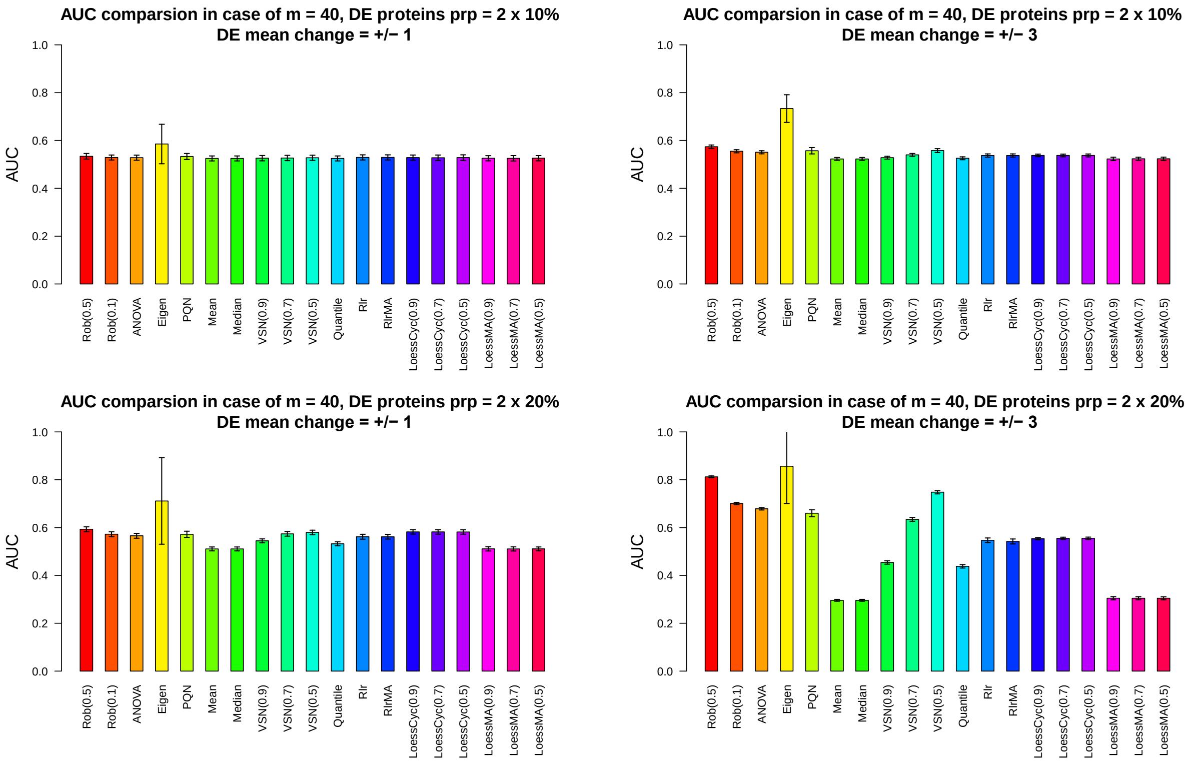


*Figure S2: AUC comparisons from various normalization methods under sample size $m = 40$ in the differential expression (DE) analysis in four situations (1) DE protein proportion = $2 \times 10\%$ and DE mean change $= \pm 1$ (the topleft panel), (2) DE protein proportion = $2 \times 10\%$ and DE mean change $= \pm 3$ (the topright panel), (3) DE protein proportion = $2 \times 20\%$ and DE mean change $= \pm 1$ (the bottomleft panel), (4) DE protein proportion = $2 \times 20\%$ and DE mean change $= \pm 3$ (the bottomright panel). The height of each bar is the mean AUC from 20 repeated simulations. The small error bars indicate one standard deviation.*

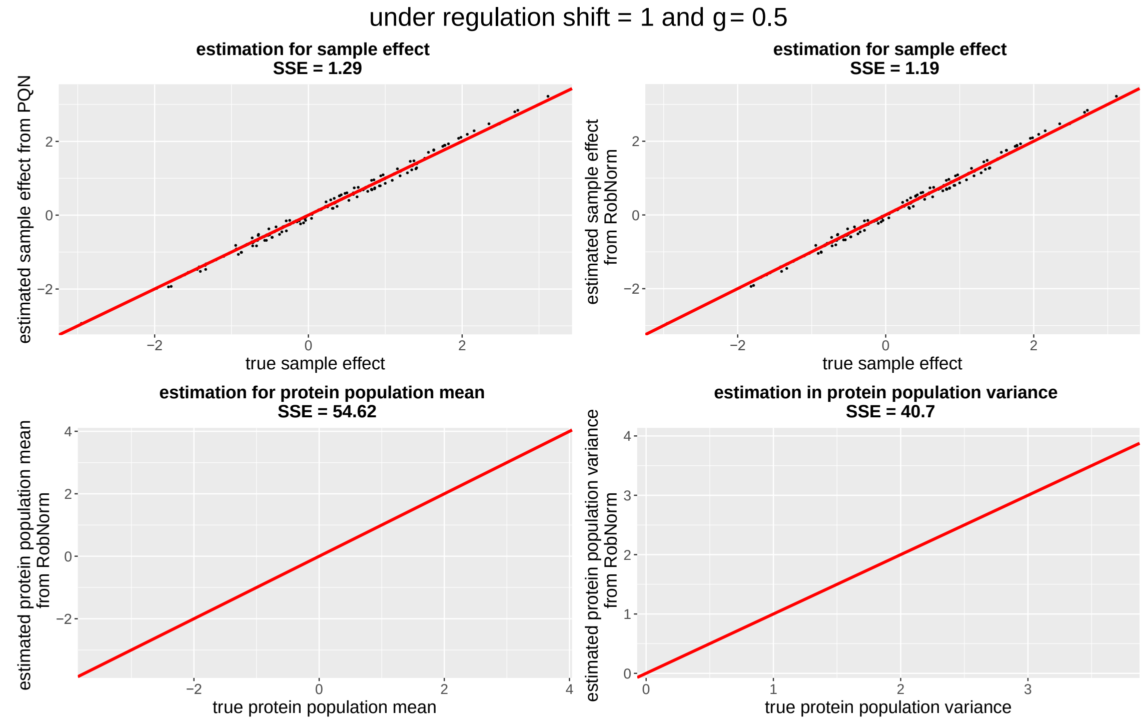


*Figure S3: Estimation performance of PQN and RobNorm under $\gamma = 0.5$ in the case of small regulation and regulation proportion as $2 \times 0.2$ in one simulated data where the data is generated from the model (1). The x-axis is for the true underlying parameter and the y-axis is for the estimate. The left upper panel is for the estimated sample effect from PQN. The rest three panels are for the estimated the sample effect from RobNorm (in the right upper panel), the protein population mean (in the left lower panel), and the protein population variance (in the right lower panel). The red line in each panel is the identity line. The sum of squared error (SSE) in summarized in each panel.*

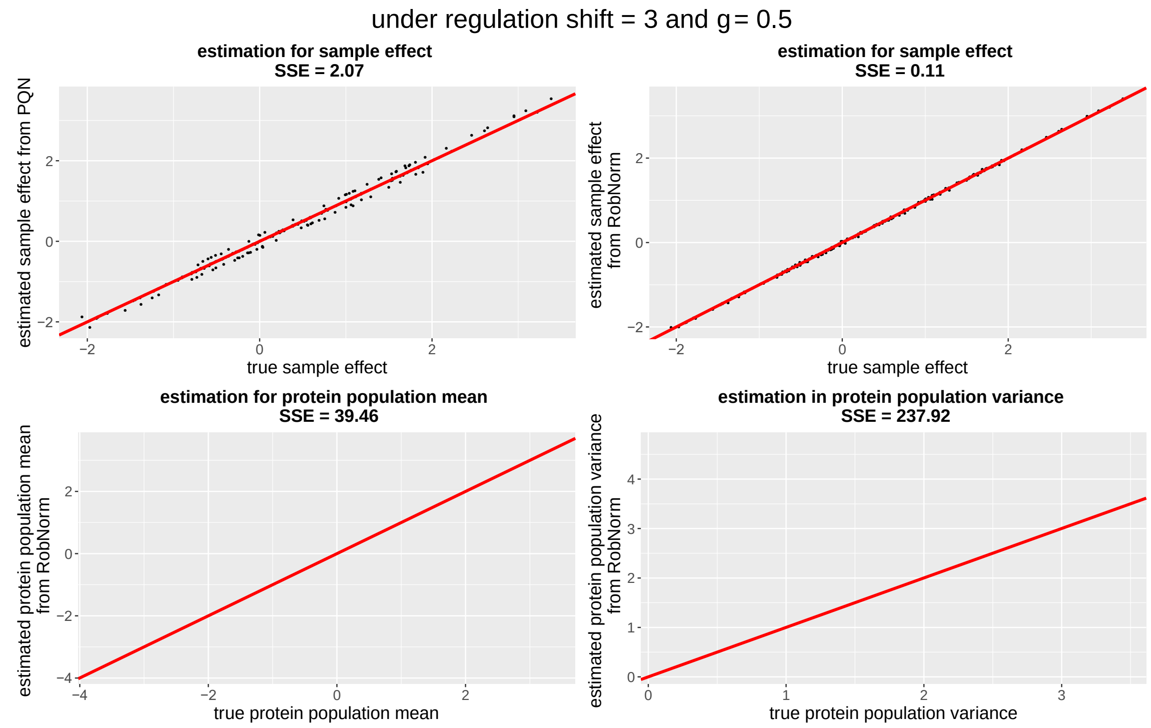


*Figure S4: Estimation performance of PQN and RobNorm under $\gamma = 0.5$ in the case of large regulation and regulation proportion as $2 \times 0.2$ in one simulated data where the data is generated from the model (1). The labels are the same as in Figure 4.*

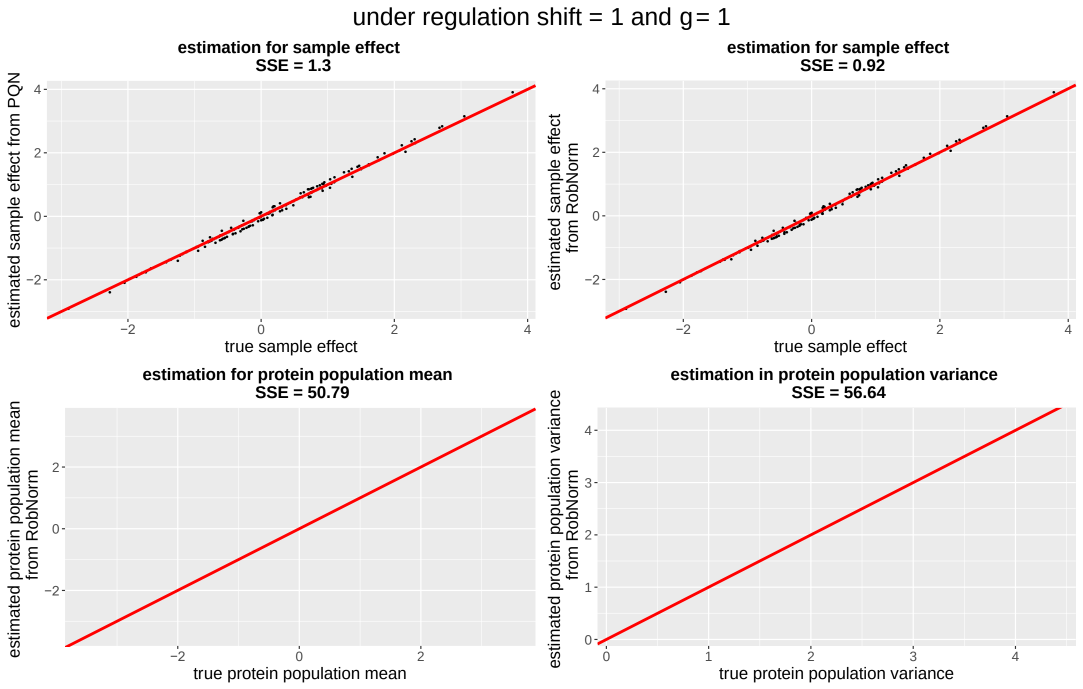


*Figure S5: Estimation performance of PQN and RobNorm under $\gamma = 1$ in the case of small regulation and regulation proportion as $2 \times 0.2$ in one simulated data where the data is generated from the model (1). The labels are the same as in Figure 4.*

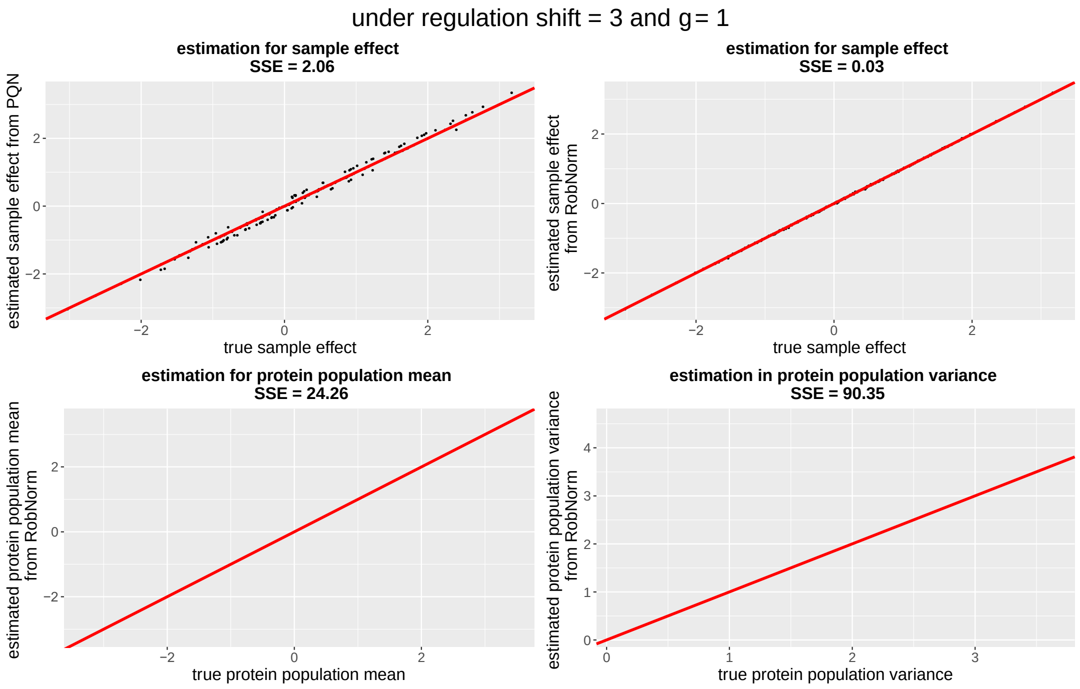


*Figure S6: Estimation performance of PQN and RobNorm under $\gamma = 1$ in the case of large regulation and regulation proportion as $2 \times 0.2$ in one simulated data where the data is generated from the model (1). The labels are the same as in Figure 4.*

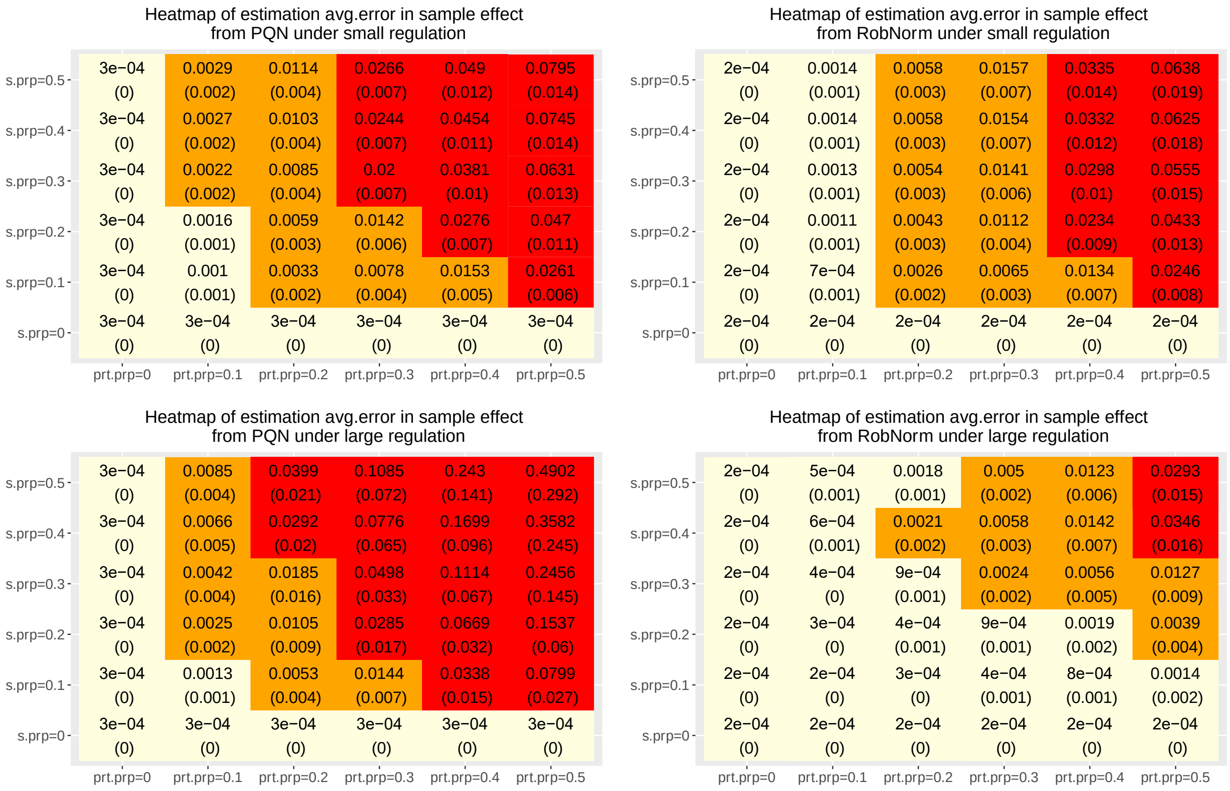


*Figure S7: Estimation error comparisons of RobNorm to PQN in estimating the sample effect* $\boldsymbol{\nu}$ *under various sizes of affected protein proportions* ($prt.prp$) *and sample proportions* ($s.prp$) *in the cases of small regulation with* $\Delta\mu = 1$ *(the panels in the first row) and large regulation with* $\Delta\mu = 3$ *(the panels in the second row). The simulated data is generated from model (1) under* $n = 5000$, $m = 200$, *and* $\gamma = 0.5$ *with various* ($prt.prp$, $s.prp$) *in the grid of* $(0.1, 0.2, 0.3, 0.4, 0.5) \times (0.1, 0.2, 0.3, 0.4, 0.5)$. *Each cell in the heatmap summarizes the estimation error in each setting. The first row in one cell is the average of the estimation error defined in (2) from 50 replicates and its second row is its standard deviation. The yellow region represents the average error* $< 10/n = 0.002$; *the orange region represents the average errors* $\geq 10/n$ *but* $< 100/n = 0.02$; *the red region represents the average error* $\geq 100/n$.

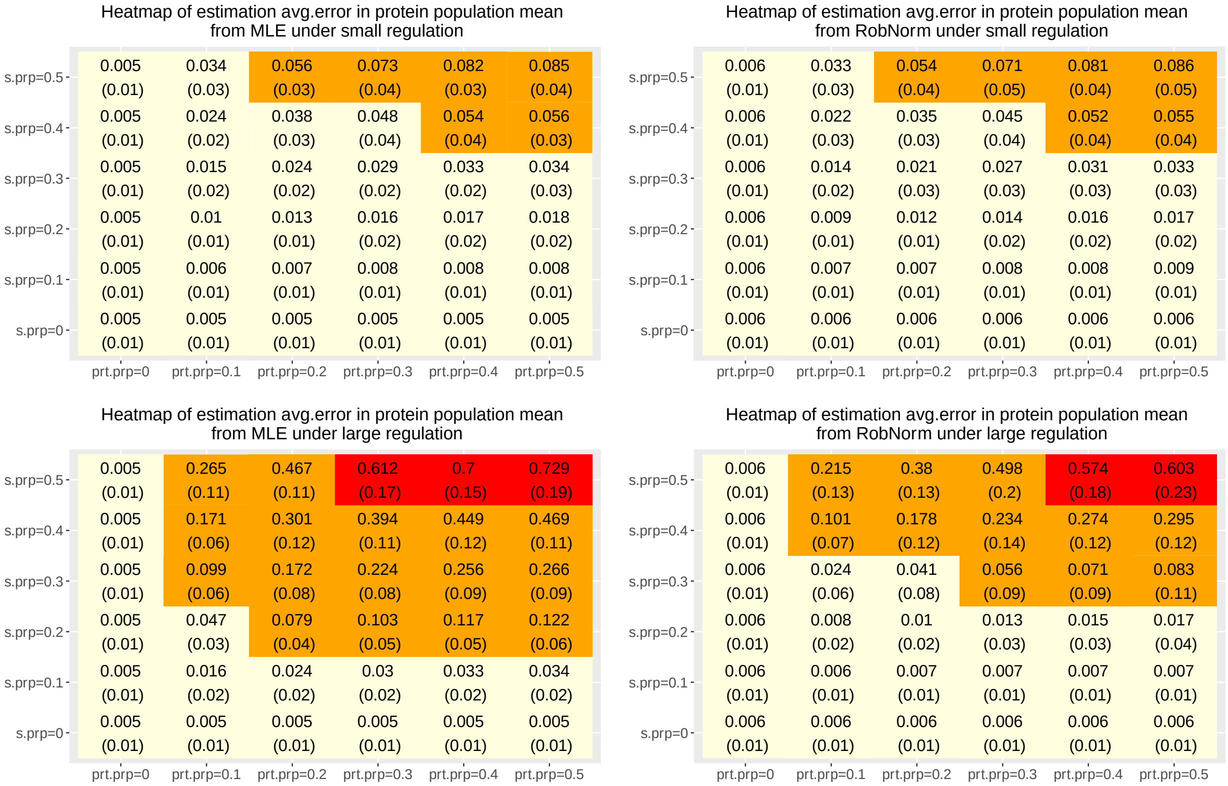


*Figure S8: Estimation error comparisons of RobNorm to the MLEs in the protein population mean $\boldsymbol{\mu}_0$ under various sizes of affected protein proportions ($prt.prp$) and sample proportions ($s.prp$) in the cases of small regulation with $\Delta\mu = 1$ (the panels in the first row) and large regulation with $\Delta\mu = 3$ (the panels in the second row). The simulated data is generated from model (1) under $n = 5000$, $m = 200$, and $\gamma = 0.5$ with various ($prt.prp$, $s.prp$) in the grid of $(0.1, 0.2, 0.3, 0.4, 0.5) \times (0.1, 0.2, 0.3, 0.4, 0.5)$. Each cell in the heatmap summarizes the estimation error in each setting. The first row in one cell is the average of the estimation error defined in (3) from 50 replicates and its second row is its standard deviation. The yellow region represents the average errors $< 10/m = 0.05$; the orange region represents the average errors $\geq 10/m$ but $< 100/m = 0.5$; the red region represents the average errors $\geq 100/m$.*

#### Real data application

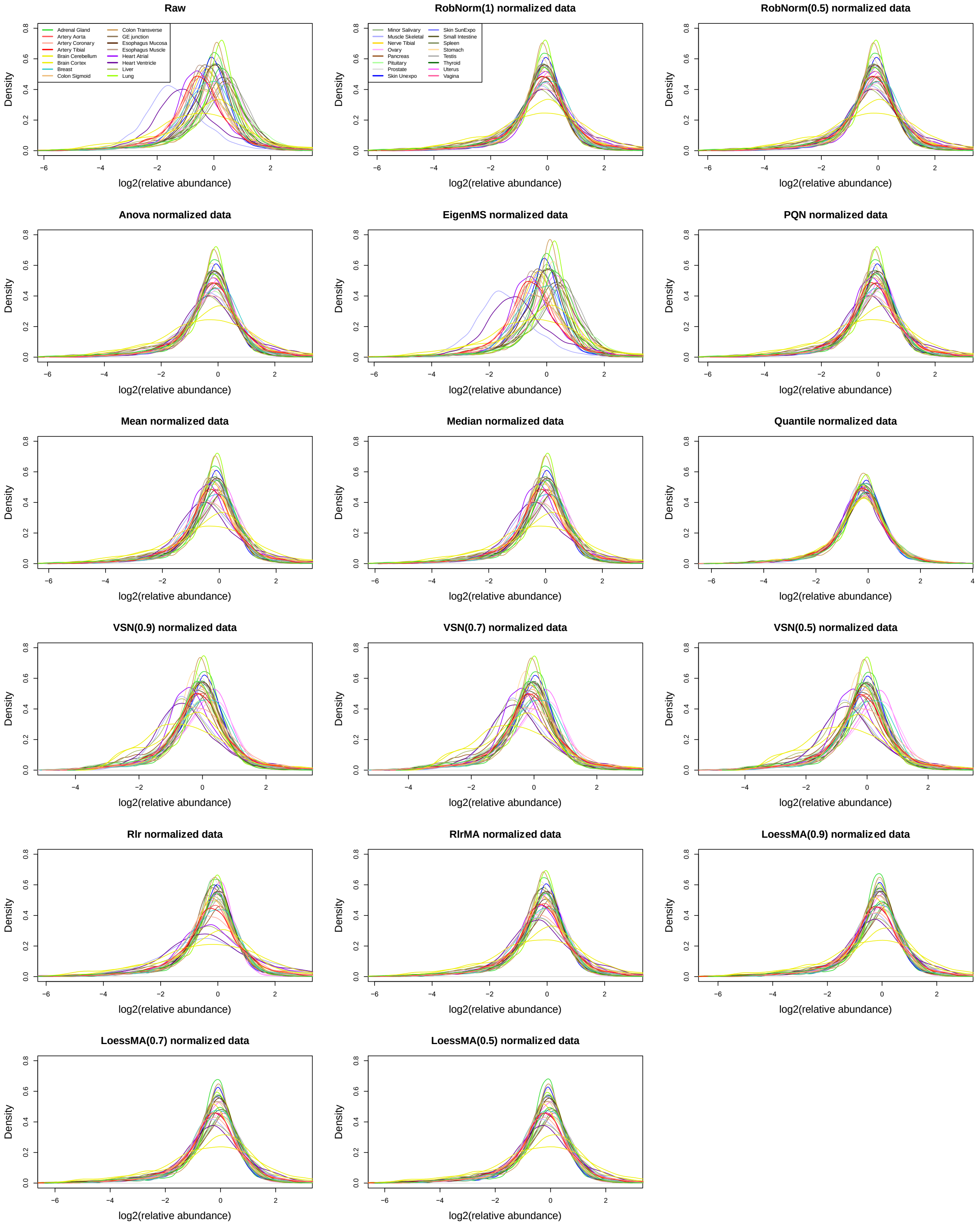


*Figure S9: The distribution of relative protein abundances in log scale at base 2 across tissues. The protein expression from each tissue is summarized from sample medians in the same tissue. The legends for the tissue types are divided in the first two panels.*

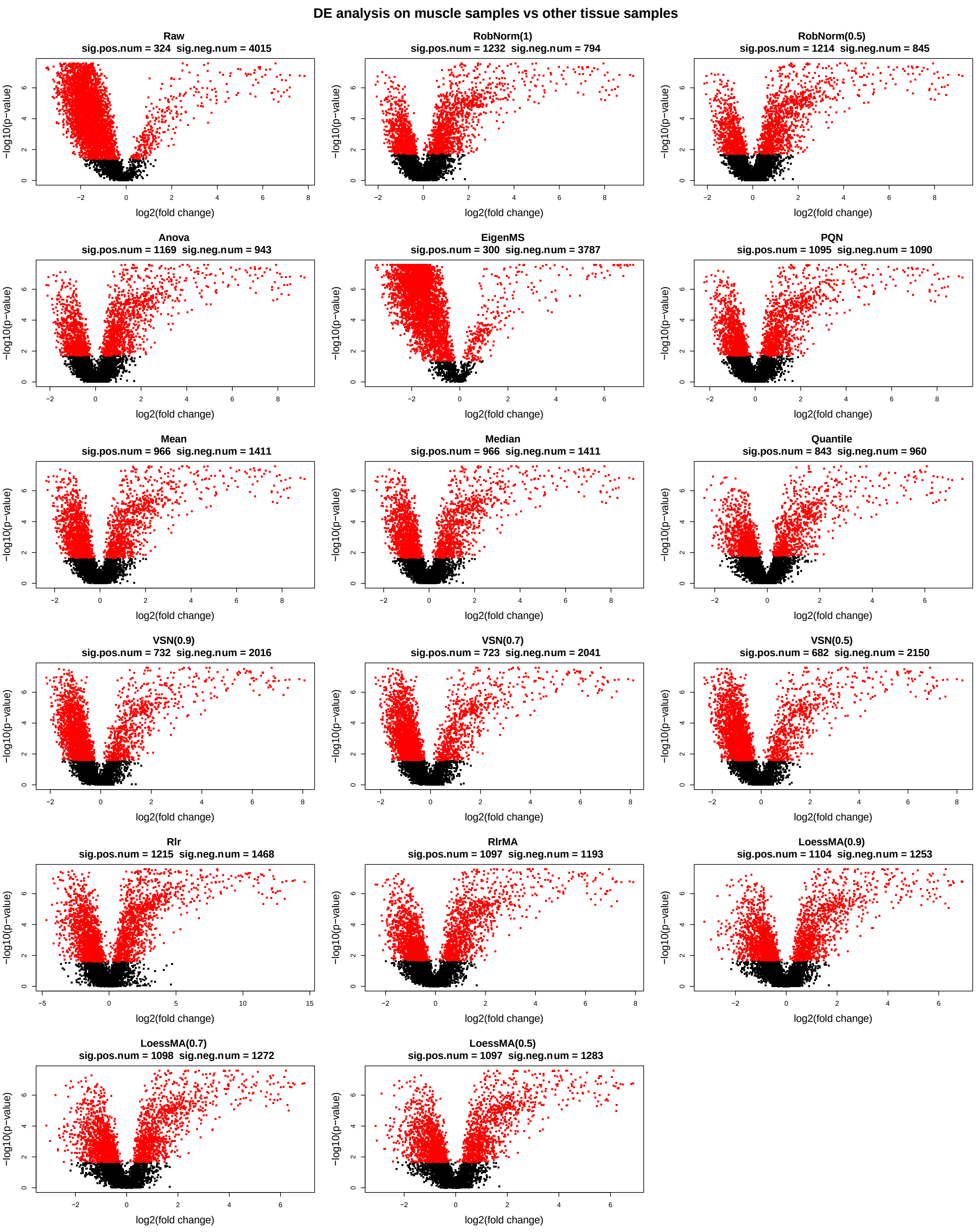


*Figure S10: Volcano plot from the differential expression (DE) analysis for muscle sample expression vs non-muscle sample expression. The p-value is obtained from the Wilcoxon rank sum test. The red dots indicate significant proteins under BH adjusted p-value < 0.05 (Benjamini and Hochberg, 1995). In the subtitle of each panel, number of significantly highly-expressed proteins (sig.pos.num) and number of significantly lowly-expressed proteins (sig.neg.num) are reported.*
